# A Genome-Wide Association Analysis Reveals Epistatic Cancellation of Additive Genetic Variance for Root Length in *Arabidopsis thaliana*

**DOI:** 10.1101/008789

**Authors:** Jennifer Lachowiec, Xia Shen, Christine Queitsch, Örjan Carlborg

## Abstract

Efforts to identify loci underlying complex traits generally assume that most genetic variance is additive. Here, we examined the genetics of *Arabidopsis thaliana* root length and found that the narrow-sense heritability for this trait was statistically zero. This low additive genetic variance likely explains why no associations to root length could be found using standard additive-model-based genome-wide association (GWA) approaches. However, the broad-sense heritability for root length was significantly larger, and we therefore also performed an epistatic GWA analysis to map loci contributing to the epistatic genetic variance. This analysis revealed four interacting pairs involving seven chromosomal loci that passed a standard multiple-testing corrected significance threshold. Explorations of the genotype-phenotype maps for these pairs revealed that the detected epistasis cancelled out the additive genetic variance, explaining why these loci were not detected in the additive GWA analysis. Small population sizes, such as in our experiment, increase the risk of identifying false epistatic interactions due to testing for associations with very large numbers of multi-marker genotypes in few phenotyped individuals. Therefore, we estimated the false-positive risk using a new statistical approach that suggested half of the associated pairs to be true positive associations. Our experimental evaluation of candidate genes within the seven associated loci suggests that this estimate is conservative; we identified functional candidate genes that affected root development in four loci that were part of three of the pairs. In summary, statistical epistatic analyses were found to be indispensable for confirming known, and identifying several new, functional candidate genes for root length using a population of wild-collected *A. thaliana* accessions. We also illustrated how epistatic cancellation of the additive genetic variance resulted in an insignificant narrow-sense, but significant broad-sense heritability that could be dissected into the contributions of several individual loci using a combination of careful statistical epistatic analyses and functional genetic experiments.

**Author summary:** Complex traits, such as many human diseases or climate adaptation and production traits in crops, arise through the action and interaction of many genes and environmental factors. Classic approaches to identify contributing genes generally assume that these factors contribute mainly additive genetic variance. Recent methods, such as genome-wide association studies, often adhere to this additive genetics paradigm. However, additive models of complex traits do not reflect that genes can also contribute with non-additive genetic variance. In this study, we use *Arabidopsis thaliana* to determine the additive and non-additive genetic contributions to the phenotypic variation in root length. Surprisingly, much of the observed phenotypic variation in root length across genetically divergent strains was explained by epistasis. We mapped seven loci contributing to the epistatic genetic variance and validated four genes in these loci with mutant analysis. For three of these genes, this is their first implication in root development. Together, our results emphasize the importance of considering both non-additive and additive genetic variance when dissecting complex trait variation, in order not to lose sensitivity in genetic analyses.

## Introduction

Identifying the loci underlying quantitative phenotypes is among the central challenges in genetics. The current quantitative genetics paradigm is based on the assumption of additive gene action, despite an increasing body of evidence showing that non-additive effects are crucial to many, if not most, biological systems [1–4]. The key arguments for remaining within the additive paradigm are that many genetic architectures with non-additive gene action display considerable additive genetic variance in populations [5] and that additive model-based approaches have facilitated detection of thousands of loci associated with many complex traits [6]. However, the amount of additive variance contributed by epistasis for example, will vary with the allele-frequencies in populations and therefore be a dynamic property of studied populations [7–13 and references therein]. Therefore, the focus on additive variation alone can leave a considerable amount of genetic variance unexplored [3, 14], and for a full dissection of a complex trait, it is often necessary to explore alternative approaches that capture non-additive variation. Statistical epistasis has been shown to be pervasive in both humans and model organisms [2, 14]. Epistatic contributions to complex traits have primarily been identified in experimental populations using candidate gene approaches [1, 2, 4, 15–19] and QTL mapping, for example [1, 20]. Genome-wide association (GWA) analyses have more recently emerged as an effective method for mapping loci underlying phenotypic variation for complex traits also in natural populations. However, GWA analyses to detect epistatic interactions are still under-utilized in efforts to identify loci contributing to the genetic architecture of complex traits. To obtain a more complete compilation of the loci contributing to the variation of a complex trait, it would therefore be valuable to also account for epistatic interactions in addition to additive effects in existing and future datasets.

The drawbacks of studying epistasis in natural populations of higher organisms include the cost and time to collect large datasets, the lack of power due to noisy phenotypes, and the difficulties in performing the necessary follow-up studies to replicate the epistatic associations, identify the polymorphic genes, and dissect the underlying mechanisms leading to statistical epistatic associations. Efforts to utilize epistatic interactions for detection of novel loci, however, are of importance for enhancing sensitivity in statistical analyses [21], for selecting the most promising analytical approaches and experimental designs for future work, and for providing insights as to what type of genetic and biological mechanisms can lead to non-additive genetic variance and genotype-phenotype relationships. Public resources have recently been developed for studying genotype-phenotype associations with GWA analyses in model organisms such as *Arabidopsis thaliana* [22] and *Drosophila melanogaster* [23]. Many powerful experimental approaches are available in these species, making them attractive for studies aiming to functionally dissect more complex genetic mechanisms such as those underlying statistical epistatic associations. For these reasons, these resources have the potential to contribute to the fundamental work of clarifying how statistical epistatic analyses should more generally be interpreted and used in efforts to dissect the genetic architectures of complex traits.

Two major challenges have often been discussed in relation with GWA analyses based on statistical epistatic models: the lack of sufficiently large experimental datasets to obtain reasonable statistical power and the computational complexity when screening the genome for multi-locus epistasis. One of the causes of low power is the need to use stringent, multiple-testing corrected significance-thresholds to account for the large number of statistical tests performed when exploring epistasis between all possible combinations of loci [24]. As allele-frequencies are often skewed in the populations used for GWA analyses, power is decreased further due to the small number of observations in the multi-locus, minor-allele genotype classes. To some extent these challenges can be overcome by increasing the sample size and making efficient use of high-performance computing [25]. Leveraging genotyped populations of model organisms can also overcome statistical limitations. The genomes of model organisms are often smaller, which reduces the search-space during genome-wide scans. The existence of inbred populations, such the *A. thaliana* RegMap [26] or 1001 Genomes Project [14] or the *D. melanogaster* Drosophila Genetic Reference Panel [15] collections, provide advantages in power compared to analyses in general outbred populations with heterozygotes. This is particularly true in analyses of epistasis due to the simpler genotype-phenotype maps. Several standard additive GWA analyses have been previously performed for many traits using the publically available *A. thaliana* collections and data sets [27, 28], including several studies of *A. thaliana* root phenotypes [29–31]. *A. thaliana* has a small genome (∼125 Mb), large numbers of readily available inbred accessions, a large knowledge base on how to efficiently score quantitative phenotypes in large numbers of individuals, and powerful experimental resources, such as collections of knock-out lines. This makes such populations into particularly well-suited models for performing exhaustive scans for epistasis and for evaluating whether non-additive genetic analyses are able to uncover novel contributions of genes to the studied phenotypes. Further, these advantages lead to a smaller theoretical decrease in power than expected due to sample size alone.

Model organism GWA studies, however, are often based on a hundred or fewer individuals, and valid concerns have been raised regarding the challenges of testing for associations with hundreds of thousands of markers, or millions of combinations of epistatic markers in such small samples. In particular, the chance of detecting a random association will increase especially when the number of individuals in the minor two-locus genotype class is small. This situation is not accounted for by common statistical multiple-testing correction. Further, there is also a risk of statistical epistatic associations arising due to “apparent epistasis” [32] in which the multi-locus genotype tags an unobserved, single polymorphism in the genome. Here, the genetic variance used to detect the association originates from a true genetic effect in the genome, but this effect is hidden in a standard GWA due to its low correlation with the individual genotyped markers. Instead the high-order correlation between the hidden genetic variant and a multi-marker genotype leads to an epistatic variance that can be detectable in the epistatic GWA analysis. Although this scenario has only previously been discussed for linked variants [32], it could occur for unlinked loci if the analyzed population is small and the number of evaluated combinations of loci is large. Also, this scenario goes beyond standard statistics used to account for multiple testing in GWA analyses, as the increased risk for false positives is not due to the strength of the statistical association. Instead, it results from the inability to properly compare alternative explanatory genetic models for the association using the available genotype data. Unless particular care is taken, both of these scenarios could lead to an increased false-positive rate among the reported loci, as well as incorrect interpretations of the presence of biological interactions if significant pairwise statistical epistasis is interpreted as evidence of an underlying genetic architecture with genetic interactions. These potential risks associated with epistatic GWA analyses in small experimental populations should therefore be considered when such analyses are used.

Here, we conduct a GWA study to dissect the genetics underlying *A. thaliana* root length in a population of wild-collected accessions. Root length had a statistically zero narrow-sense heritability in this population, but a larger significant broad-sense heritability. The genetic contribution to this trait was therefore unlikely to be detected using standard additive genetic models, and it was not surprising that no locus was associated with root-length in the standard additive-model based GWA analyses. An exhaustive GWA scan for two-locus interactions, however, revealed four genome-wide significant interacting pairs. All of these displayed genotype-phenotype relationships in which epistasis cancelled the marginal additive genetic effects of the interacting loci, explaining the low levels of additive genetic variance in the population. A new statistical method was used to evaluate the robustness of the GWA associations in the small population, and it was estimated that half of these pairs are at risk of being false positives. We also found low risk that the statistical epistatic interactions were caused by “apparent epistasis” [32]. Our experimental evaluations of genes in the associated regions using T-DNA insertion mutants identified functional candidates that affected root length in four regions that were part of three pairs, from which three genes were newly implicated in root development [33]. Together, our results illustrate the value of epistatic GWA analyses for revealing novel loci contributing to complex traits in small, inbred populations. Further, we noticed an increased risk for false-positive epistatic associations and “apparent epistasis” due to the small sample-size but the top-ranked associations proved, despite this, to be valuable in further experimental validation, as several functional candidate genes were identified in loci that would have been discarded based on stringent statistics alone. We also report several promising functional candidate genes for future studies to dissect the molecular mechanisms underlying root development in *natural* A. thaliana populations that would have remained undetected without an epistatic GWA analysis.

## Results

### Epistasis is the primary source of genetic variance for root length

To study the genetic regulation of mean root length, we generated a comprehensive data set containing the *A. thaliana* root mean lengths across 93 wild-collected accessions (Fig S1, Table S1). The 93 accessions used here were previously genotyped at ∼215,000 single nucleotide polymorphisms (SNPs) and used in a standard GWA analysis for many plant phenotypes [27]. We grew seedlings in a randomized design on standard plant medium in temperature-controlled chambers to reduce environmental variation among individuals [34]. This experimental design allows for parsing of genetic and environmental factors underlying root length [35, 36].

In our dataset, mean root length varied significantly across accessions (p < 1 × 10^−10^, ANOVA; Fig1). To evaluate whether the observed variation in mean root length was under genetic control, we estimated both the narrow-and broad-sense heritability. This analysis used a linear mixed model that simultaneously estimated the polygenic additive and epistatic variance components using the R/hglm package [37]. We estimated the total broad sense heritability to be 0.24 (95% CI 0.10 – 0.38) for mean root length (Table 1), showing that the trait is under genetic control. We also partitioned the genetic variance into the individual additive and epistatic variance components and found that almost all the variance was epistatic (0.22; p = 1.32 × 10^−6^) and only a small, insignificant fraction was additive (0.01; p = 0.63). These heritability patterns suggest that an epistatic GWA analysis might be needed to identify the loci explaining the heritable phenotypic variation in root length mean.

**Fig 1.**
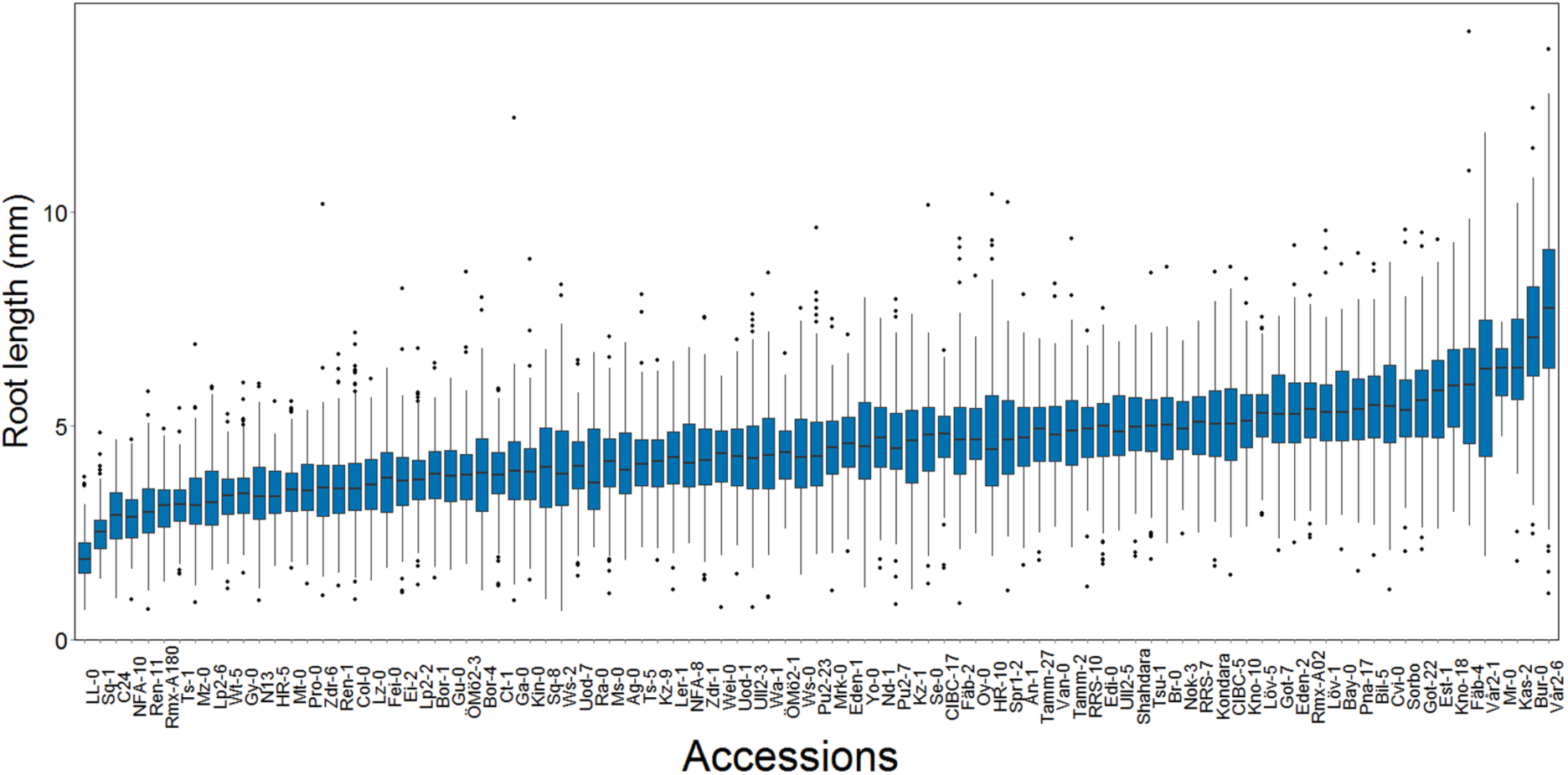
The distributions of within-accession root length mean and variance in a sample of 93 natural *A. thaliana* accessions. There is a large dispersion of the root length mean at day seven among the accessions (*n* = 3 × 70 dark grown seedlings / genotype).

**Table 1.** Estimated narrow (h^2^) and broad sense (H^2^) heritabilities of root length mean and variance in a population of 93 natural *A. thaliana* accessions. The broad-sense heritability was intermediate for root length mean, with a negligible additive contribution.

| Trait | $H^2$ | | $h^2$ |
| --- | --- | --- | --- |
|  | ANOVA<sup>1</sup> | hglm<sup>2</sup> | hglm<sup>2</sup> |
| Mean root length (mm) | 0.25 | 0.24 | 0.015 |
| | (p = 0) | (s.e. = 0.09, p = $7.2 \times 10^{-3}$)<sup>3</sup> | (s.e. = 0.056, p = 0.79)<sup>3</sup> |

<sup>1</sup>Estimated using ANOVA with accession as a fixed effect, <sup>2</sup>Estimated using the R/hglm package (hglm) by fitting a linear mixed model including both additive and epistatic kinship matrices as random effects, <sup>3</sup>p-values from Wald tests for the heritability being larger than zero, and s.e. are the standard errors estimated via jackknife resampling.

### No additive genome-wide associations were detected for mean root length

Two GWA analysis methods were used to screen for additive loci contributing to the phenotypic variation in mean root length. The first method was a standard single-locus additive mixed-model based approach as implemented in the R-package GenABEL [38] that accounted for population-structure by modelling the genomic kinships between the accessions as a random effect. Significance testing was done using a Bonferroni-corrected significance threshold. The second method was based on a whole-genome generalized ridge regression heteroscedastic effects model (HEM), in which all SNPs were included simultaneously and their contributions estimated as random effects using the R/bigRR package [39]. Here, a 5% genome-wide significance threshold was determined by permutation testing. None of the tested SNPs were significantly associated with mean root length in either of these analyses (Fig S1a, b). Given the low estimate for the narrow-sense heritability for mean root length in this population (h^2^ = 0.01), this outcome was not surprising.

### An epistatic genome-wide association analysis uncovered multiple interacting loci that contributed to mean root length

To identify loci that contributed to the epistatic genetic variance for mean root length, we performed an exhaustive two-locus SNP-by-SNP epistasis analysis (PLINKv1.07) [40]. Despite the limited population size, several factors increased our chances of finding novel loci using this approach: the small additive and large epistatic genetic variance (Table 1), the high precision in estimating the phenotypic values due to extensive replication, the presence of only four two-locus genotypes in the population of inbred accessions, and a reduced number of pairwise combinations to test in the small genome. To further enhance power and decrease the risk of false-positives, we excluded pairs of SNPs where the minor two-locus genotype class contained few observations. This was achieved by applying two data filters: first, only SNPs with a minor allele frequency greater than 0.25 were included, and second all SNP-by-SNP combinations with fewer than four accessions in the minor genotype class were removed. In the epistatic analyses, we accounted for population structure by performing the GWA analysis using a linear mixed model including the genomic kinship matrix. Significance for the pairs were determined using a standard Bonferroni-correction for the number of independent pair-wise tests across the independent linkage blocks across the *A. thaliana* genome [41]. In total, the epistatic analysis included approximately 78 million such independent tests. Six SNP-pairs, representing seven genomic locations and four unique combinations of loci, passed this threshold (Table 2). Two pairs were detected twice by associations to tightly linked SNPs located 1.5 kb and 505 bp apart (Chromosome 3 at 9,272,294 and 9,273,674 bp - SNPs 3_9272294/ 3_9273674 - and Chromosome at 15,862,026 and 15,862,525 bp - SNPs 5_15862026/5_15862525, respectively), and one SNP (Chromosome 3 at 66,596 bp - SNP 3_66596) was part of two unique interacting pairs (Table 2). The other two SNP-pairs contained independent SNPs (R^2^ < 0.8).

**Table 2.** Exhaustive two-dimensional GWAS-scan for epistasis identifies six significant interactions, representing four unique pairs, associated with the root length mean.

| Pair | SNP1<sup>1</sup> | SNP2<sup>1</sup> | Interaction<br>(mm) | p-value<sup>2</sup><br>(GWA) | p-value<sup>3</sup><br>(SS) |
| --- | --- | --- | --- | --- | --- |
| 1 | rs346846662<br>(3_66596) | rs347034954/rs347048583<br>(3_9273674/3_9272294) | 0.55 | $1.5 \times 10^{-10}$ | 0.003 |
| 2 | rs347005428<br>(1_17257526) | rs346595296/rs346511785<br>(5_15862026/5_15862525) | 0.63 | $1.7 \times 10^{-10}$ | 0.28 |
| 3 | rs346846662<br>(3_66596) | rs347177728<br>(5_18241640) | 0.63 | $3.0 \times 10^{-10}$ | 0.37 |
| 4 | rs347374065<br>(3_10891195) | rs347377866<br>(5_1027939) | 0.63 | $2.3 \times 10^{-10}$ | 0.95 |

<sup>1</sup>Name of associated SNP(s) at the epistatic locus as on the Affymetrix single nucleotide polymorphism (SNP) chip (our abbreviation as Chromosome\_PositionInBp); <sup>2</sup>p-value for the interaction effect of the two loci in the epistatic GWA analysis [40]; <sup>3</sup>Probability that the pair is a false-positive when accounting for the small sample-size (SS) [42].

### Explorations of epistatic genotype-phenotype maps revealed an epistatic cancellation of the additive genetic variance

The four SNP pairs that were significant in the epistatic GWA (Table 2; Fig 2a) explained no additive variance, but together 51% of the phenotypic variance of mean root length (95% CI: 31.2% - 72.4% from bootstrap). To explore the origin of the low additive variance in the presence of the revealed epistasis, we estimated the multi-locus genotype-phenotype (G-P) maps and their respective genotype frequencies in the analyzed population. The G-P maps were highly related for the four pairs: the minor allele double-homozygote was always associated with the longest mean root length and the shortest mean root lengths were associated with the genotypes combining one major-and one minor-allele homozygote (Fig 2b; Fig S2). Although this type of epistasis will lead to marginal additive genetic effects for the two loci for many allele-frequencies, there are also combinations of allele-frequencies where the epistasis will cancel the additive genetic variance completely. We illustrate this epistatic cancellation of the additive genetic variance for the G-P map and allele-frequencies of the most significant pair in Fig 2c. For this pair, there are minimal differences in the average (marginal) mean root lengths for the two genotypes at the individual loci, and hence a minor contribution to the additive genetic variance. Their joint contribution to the phenotypic variance, however, is large due to the large differences in root lengths among the four two-locus genotype classes. The epistatic GWA analysis is able to reveal these pairs in this population as the differences between these genotypes contribute to the epistatic genetic variance instead.

**Fig 2.**
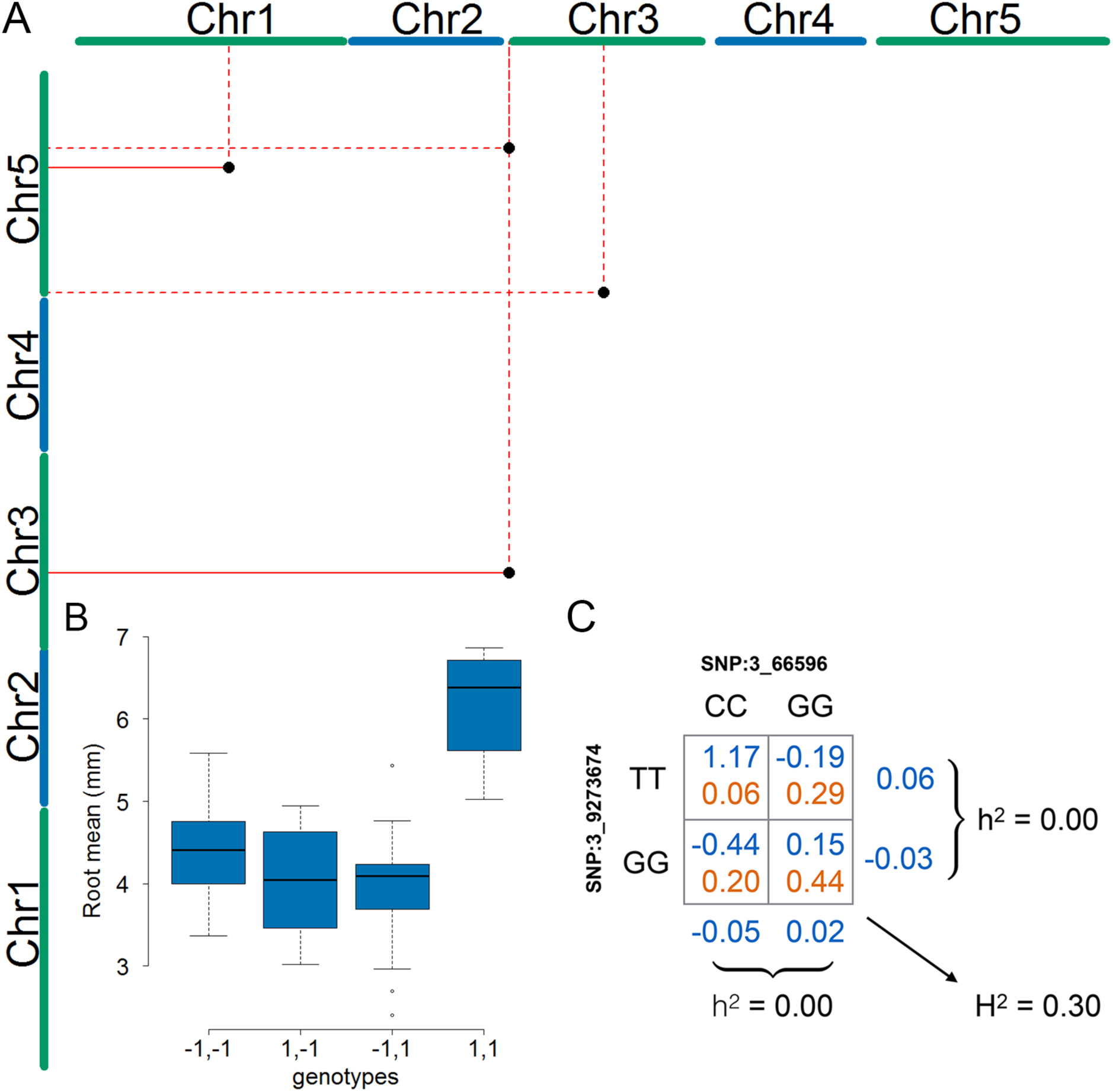
Four statistical epistatic interactions associated with mean root length. (A) The x-and y-axes represent the five *A. thaliana* chromosomes. The positions of the seven SNPs that are part of the four significant interacting pairs (Table 2) are indicated by a black dot. Solid lines indicate support for an interaction by more than one linked SNP and dotted lines indicate support by a single SNP. (B) Genotype-phenotype (G-P) map of the root means for the four genotype combinations constituting the interaction between the SNPs on chromosome 1 (17,257,526 bp) and chromosome 5 (15,862,026 bp). The major allele is indicated by -1, and the minor allele is indicated by 1. This G-P-map is a representative example for the other pairs in which the accessions with a minor and major allele have lower phenotypic values than those with either both major or both minor alleles (Fig S2). (C) G-P map for the most significant epistatic pair illustrates how this type of epistasis cancels the additive genetic variance at the allele-frequencies observed for the two loci in the analyzed population of wild-collected *A. thaliana* accessions.

### Evaluating the robustness of statistical epistatic associations to a small population size

We next applied a conservative genome-wide test to estimate the risk that the epistatic GWA associations were false positives due to the small population size. A pseudo-marker was created for each pair of SNPs in the epistatic scan, resulting in an epistatic pseudo-genome with 78 million markers. Given the epistatic effect estimated for the four pairs that reached genome-wide significance in the epistatic GWA, we computed the genome-wide probability of these pairs being false-positives using the R/p.exact package [42], assuming conservatively that all the pseudo-markers are independent [42] (Table 3). The pair 3_10891195/5_1027939 was found to be sensitive to population size (p = 0.94). The pairs 1_17257526/5_15862026 and 3_66596/5_18241640 were less sensitive (p = 0.28 and p = 0.37, respectively), and hence there is a 90% probability that at least one of these two signals is not false. The last pair 3_66596/3_9273674 was not sensitive to population-size (p = 0.003). We conclude that it is essential to consider the effect of population-size when interpreting the results from GWA analyses in small populations. In our particular case, half of the original associations were at risk of being false-positives due to small populations size.

**Table 3.** Estimation of the risk that epistatic pairs identified in the GWA analysis are due to high-order LD to unobserved functional variants (“apparent epistasis”). Using the whole-genome re-sequencing data from the reference 1001-genomes *A. thaliana* collection, we estimated the high-order LD (r^2^) between the four pseudo-markers representing our epistatic pairs and all sequencing variants that were not genotyped in our GWA analysis. A bootstrap approach was used to estimate the mean and max r^2^ between the epistatic pseudo-markers and all the genome-wide sequencing-variants. The risk that an association might be due to “apparent epistasis” was calculated as the risk of observing an r^2_max_^ > 0.8 in this analysis.

| Significant pairs in two-dimensional epistatic GWA analysis |  |  |  |  |
| --- | --- | --- | --- | --- |
| Locus 1<sup>1</sup> | 3_66596 |  | 1_17257526 | 3_10891195 |
| Locus 2<sup>1</sup> | 3_9273674 | 5_18241640 | 5_15862026 | 5_1027939 |
| $r_{\text{mean}}^2$ | 0.01 ± 0.001 | 0.01 ± 0.001 | 0.01 ± 0.003 | 0.01 ± 0.001 |
| $r_{\text{max}}^2$ | 0.52 ± 0.10 | 0.66 ± 0.15 | 0.78 ± 0.11 | 0.78 ± 0.09 |
| $P(r_{\text{max}}^2 > 0.8)$ | 0.002 | 0.18 | 0.42 | 0.42 |
| MAF<sub>PM</sub> | 0.065 | 0.043 | 0.043 | 0.043 |

<sup>1</sup>Locus detected as part of epistatic pair named as Chromosome\_PositionInBp ; $r_{\text{mean}}^2$ : mean high-order LD between epistatic pseudo-marker and genome-wide sequence variants ± standard error; $r_{\text{max}}^2$ : maximum high-order LD between epistatic pseudo-marker and genome-wide sequence variants ± standard error; $P(r_{\text{max}}^2 > 0.8)$ : probability of observing a high-order LD larger then 0.8 in a random sample of 93 accessions with MAF<sub>PM</sub> for the epistatic pseudo-marker; MAF<sub>PM</sub>: minor-allele frequency for the epistatic pseudo-marker, i.e. the frequency of the minor-allele double-homozygote for the epistatic pair.

### Quantifying the risk of “apparent epistasis” for significant statistical epistatic associations

For “apparent epistasis” to be present, the hidden functional variant needs to be in high linkage disequilibrium (measured as r^2^) with one of the “epistatic pseudo-markers” described above. Therefore, “apparent epistasis” involving physically unlinked markers is only expected for very strong mutations. However, if any such hidden variant exists, by definition neither its genetic effect nor its linkage disequilibrium to the “epistatic pseudo-markers” can be estimated from the GWA datasets. However, if a GWA study is performed in a species with sequenced reference populations, the genomic data from that population can be used to estimate the risk of observing a high r^2^ between the tested markers and other polymorphisms in the genome. Here, we used the 1001 Genomes Project *A. thaliana* genomes [22] to perform this analysis for the four significant pairs in the epistatic GWA analysis. We defined bi-allelic “epistatic pseudo-markers” by assigning individuals with the two-locus minor-allele homozygote genotype the minor pseudo-marker genotype and all others the major pseudo-marker genotype. By using a bootstrap analysis based on samples of 93 accessions from the 728 whole-genome re-sequenced accessions from the 1001 Genomes Project data, we estimated the probability of observing a high r^2^ (r^2^>0.8) between any individual SNP in the re-sequencing data and the “epistatic pseudo-markers” for the four pairs. This analysis revealed that the “epistatic pseudo-markers” corresponding to the identified pairs have very low r^2^ to most of the hidden variants (mean r^2^ =0.01). There is a very low risk to observe a high r^2^ (r^2^ > 0.8) to any hidden variant in our population for the most strongly associated pair (p < 0.005). For the other three pairs, this risk is still low (p = 0.18 - 0.42) but sufficiently large to indicate that the results should be interpreted with caution (Table 3).

### Functional exploration of candidate genes in the associated regions using T-DNA insertion lines

Our statistical analyses suggest that half of the pairs with genome-wide significant statistical epistasis might be false-positives due to the small population-size. As only four pairs were found, we decided to also perform an initial experimental and bioinformatics analysis that included all these pairs to evaluate whether the statistical risk of false-positives was reflected also by the presence of known, or novel, functional candidate genes for root-length in the associated regions.

We therefore first extracted the genotypes for 13 million genome-wide SNPs from the whole-genome re-sequencing data from the 1001 Genomes Project [43]. SNPs that were in LD (r^2^ > 0.8) and located within 5kb upstream or downstream of the leading SNP in the epistatic GWA were considered further as candidate functional mutations. In Table 4, we list the predicted functional effects of the SNPs that fulfill these criteria, together with their locations, for each pair. In total, thirteen genes and two transposons were found in LD with the leading SNPs in the seven regions detected in the epistatic GWA analysis to root length.

**Table 4.**
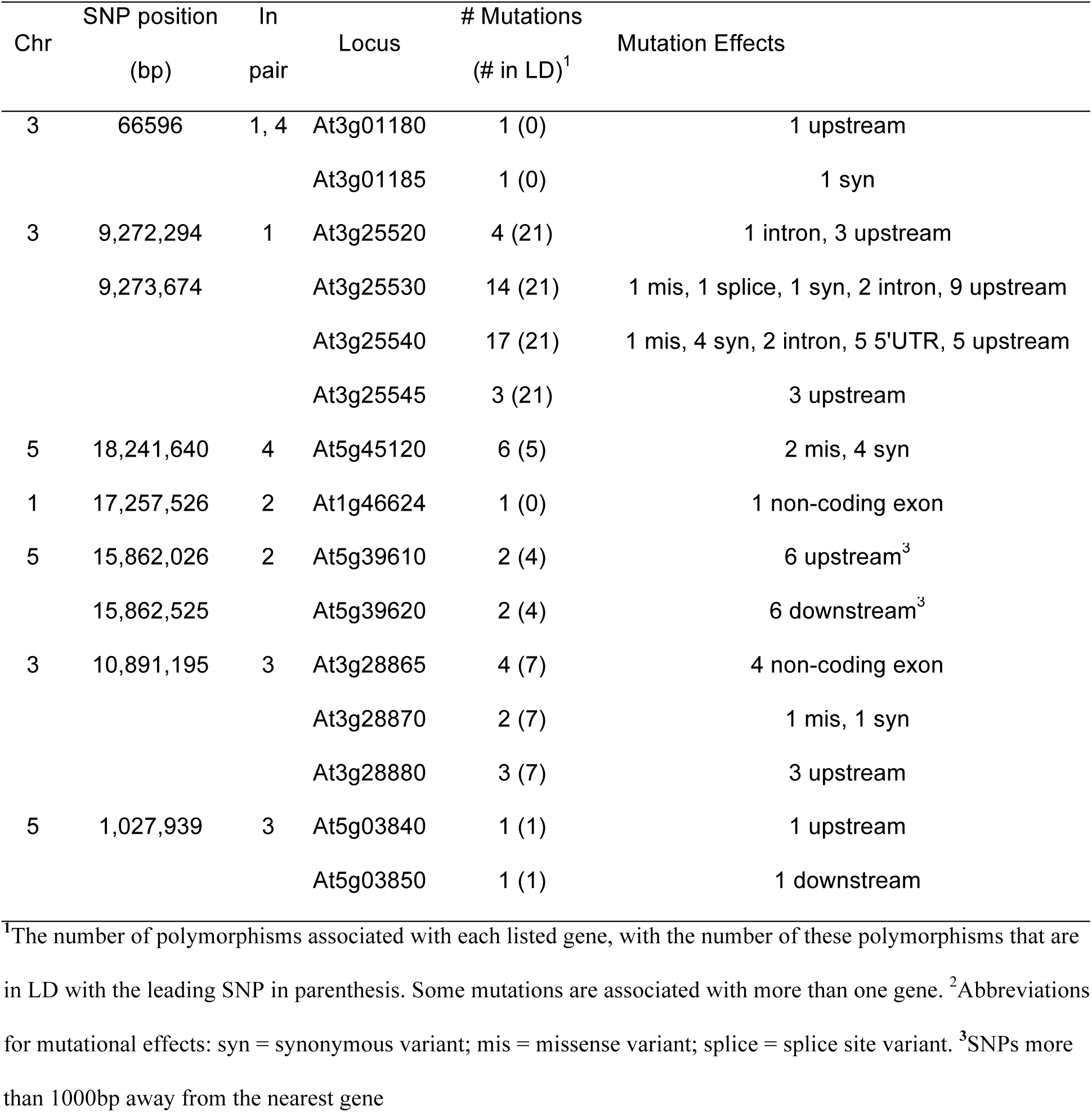
Summary of identified polymorphisms in LD with the leading SNPs from the epistatic GWA analysis. Four unique interacting pairs of loci were significantly associated with root length (Table 2). Here, the genes located within 1000bp of the leading SNPs, together with the mutational effects of the identified polymorphisms, are listed.

| Chr | SNP position (bp) | In pair | Locus | # Mutations (# in LD)<sup>1</sup> | Mutation Effects |
| --- | --- | --- | --- | --- | --- |
| 3 | 66596 | 1, 4 | At3g01180 | 1 (0) | 1 upstream |
|  |  |  | At3g01185 | 1 (0) | 1 syn |
| 3 | 9,272,294 | 1 | At3g25520 | 4 (21) | 1 intron, 3 upstream |
|  | 9,273,674 |  | At3g25530 | 14 (21) | 1 mis, 1 splice, 1 syn, 2 intron, 9 upstream |
|  |  |  | At3g25540 | 17 (21) | 1 mis, 4 syn, 2 intron, 5 5'UTR, 5 upstream |
|  |  |  | At3g25545 | 3 (21) | 3 upstream |
| 5 | 18,241,640 | 4 | At5g45120 | 6 (5) | 2 mis, 4 syn |
| 1 | 17,257,526 | 2 | At1g46624 | 1 (0) | 1 non-coding exon |
| 5 | 15,862,026 | 2 | At5g39610 | 2 (4) | 6 upstream<sup>3</sup> |
|  | 15,862,525 |  | At5g39620 | 2 (4) | 6 downstream<sup>3</sup> |
| 3 | 10,891,195 | 3 | At3g28865 | 4 (7) | 4 non-coding exon |
|  |  |  | At3g28870 | 2 (7) | 1 mis, 1 syn |
|  |  |  | At3g28880 | 3 (7) | 3 upstream |
| 5 | 1,027,939 | 3 | At5g03840 | 1 (1) | 1 upstream |
|  |  |  | At5g03850 | 1 (1) | 1 downstream |

<sup>1</sup>The number of polymorphisms associated with each listed gene, with the number of these polymorphisms that are in LD with the leading SNP in parenthesis. Some mutations are associated with more than one gene. <sup>2</sup>Abbreviations for mutational effects: syn = synonymous variant; mis = missense variant; splice = splice site variant. <sup>3</sup>SNPs more than 1000bp away from the nearest gene

Further, we identified T-DNA insertion lines for 12/15 genes and transposons comprising the seven statistically epistatic loci and experimentally evaluated them for effects on root length. In Table 5, we present the genes for which the T-DNA insertions significantly altered the root length. Results for all evaluated T-DNA lines are presented in the Supplementary Text. For three of these loci (*NAC6*, *TFL1*, *At2g28880*), we are the first to show that these underlying genes contribute to root growth, and for one locus (*ATL5*), our results confirm previous findings.

**Table 5.** Function and expression patterns of genes in LD with leading SNP from epistatic GWAS analysis. The gene names or proposed function is listed for all genes harboring polymorphisms in LD with the leading epistatic SNPs in the whole genome interaction analysis.

| Chr | Position (bp) | In pair | Locus | Gene/Putative function | Highest expression tissue<sup>1</sup> |
| --- | --- | --- | --- | --- | --- |
| 3 | 66596 | 1, 4 | At3g01180 | SS2 | cauline leaves |
|  |  |  | At3g01185 | unknown | NA |
| 3 | 9,272,294/ | 1 | At3g25520 | ATL5 | apical meristem, seedling root |
|  | 9,273,674 |  | At3g25530 | LOH1 | rosette leaves |
|  |  |  | At3g25540 | GR1 | seedling root, imbibed seed |
|  |  |  | At3g25545 | unknown | seedling root (vasculature), imbibed seed |
| 5 | 18,241,640 | 4 | At5g45120 | eukaryotic aspartyl protease family protein | adult root (meristem), imbibed seed |
| 1 | 17,257,526 | 2 | At1g46624 | gypsy-like retrotransposon | NA |
| 5 | 15,862,026/ | 2 | At5g39610 | NAC6 | seedling root (vasculature) |
|  | 15,862,525 |  | At5g39620 | RAB GTPase homolog G1 | adult root vasculature, mature seed |
| 3 | 10,891,195 | 3 | At3g28865 | LINE retrotransposon | NA |
|  |  |  | At3g28870 | histone deacetylase | mature seed |
|  |  |  | At3g28880 | Ankyrin repeat family protein | NA |
| 5 | 1,027,939 | 3 | At5g03840 | TFL1 | adult root vasculature (mature zone) |
|  |  |  | At5g03850 | nucleic acid-binding, OB-fold-like protein | seedling root, apical meristem |

<sup>1</sup> Locations of high expression were obtained from the BAR eFP *Arabidopsis* Browser (Winter et al. 2007).

### T-DNA mutant analysis reveals four functional candidate genes for root development in regions detected using an epistatic genome-wide association analysis

#### *Decreased root length in* ATL5/OLI5/PGY3 *T-DNA insertion mutants*

Two linked SNPs on chromosome 3 (3_9272294 and 3_9273674; Table 2) were associated with *A. thaliana* root length through an epistatic interaction with a distal SNP on the same chromosome (3_66596). Twenty-one SNPs located in four genes were in high LD with the two linked SNPs (Table 4; Table 5). For three of these (At3g25520, At3g25530 and At3g25540), we were able to obtain insertion mutants and scored their mean root length compared to wild-type (*Col-0*) controls. The insertion mutant for At3g25520 had significantly shorter roots than wild type (p = 1.2 × 10^−4^; Tukey’s post-hoc test); the insertion mutants for At3g25530 and At3g25540 did not differ significantly (p = 1.00 and p = 0.23, respectively) from wild type (Fig 3).

**Fig 3.**
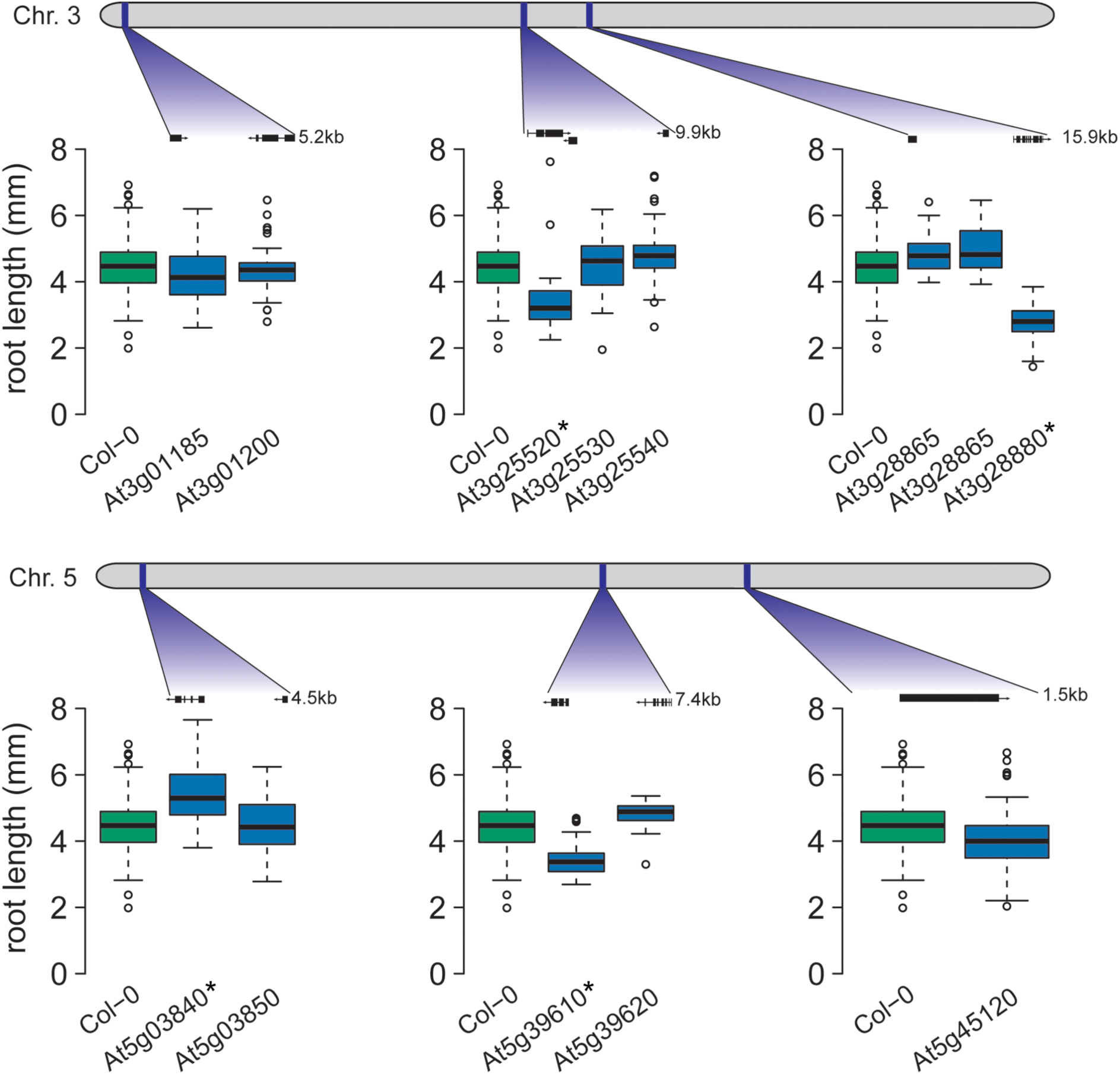
Four candidate genes exhibit altered root growth in T-DNA insertion lines. T-DNA lines for candidate genes in six of the seven epistatic regions were tested for root length differences to wild-type (*Col-0*). The gene models for the tested T-DNA insertion lines are pictured. For four of the six loci, a T-DNA line exhibited significantly different root length from *Col-0* (significantly different lines are marked with a * in the figure).

At3g25520 encodes ATL5/OLI5/PGY3, a 5S rRNA binding protein whose promoter is highly accessible in seedling roots [44], corresponding to expression in roots [45]. Our finding agrees with earlier studies, in which the loss-of-function mutant *oli5-1* exhibits shorter primary roots than wild-type [33]. In the tested accessions, three SNPs were found in the *ATL5* promoter and a fourth in an intron, suggesting variable transcriptional regulation across accessions.

#### A novel role for NAC6 in the regulation of root length

Two linked SNPs on chromosome 5 (5_15862026 and 5_15862525; Table 2) were detected by their statistical epistatic interaction with an SNP on chromosome 1 (17,257,526 bp - SNP 1_17257526; Table 2). Both of these linked SNPs, as well as the other SNPs in high LD, were intergenic with the nearest gene, *NAC6* (At5g39610; Table 5), which is located 1.9 kb away. *NAC6* is a transcription factor regulating leaf senescence [46] and is highly expressed in senescing leaves and in maturing seeds [45]. We phenotyped the available At5g39610 insertion mutant for *NAC6* and found a significant decrease in mean root length (p = 5.82 × 10^−10^; Tukey’s post-hoc test; Fig 4), strongly implicating *NAC6* in the regulation of root length.

**Fig 4.**
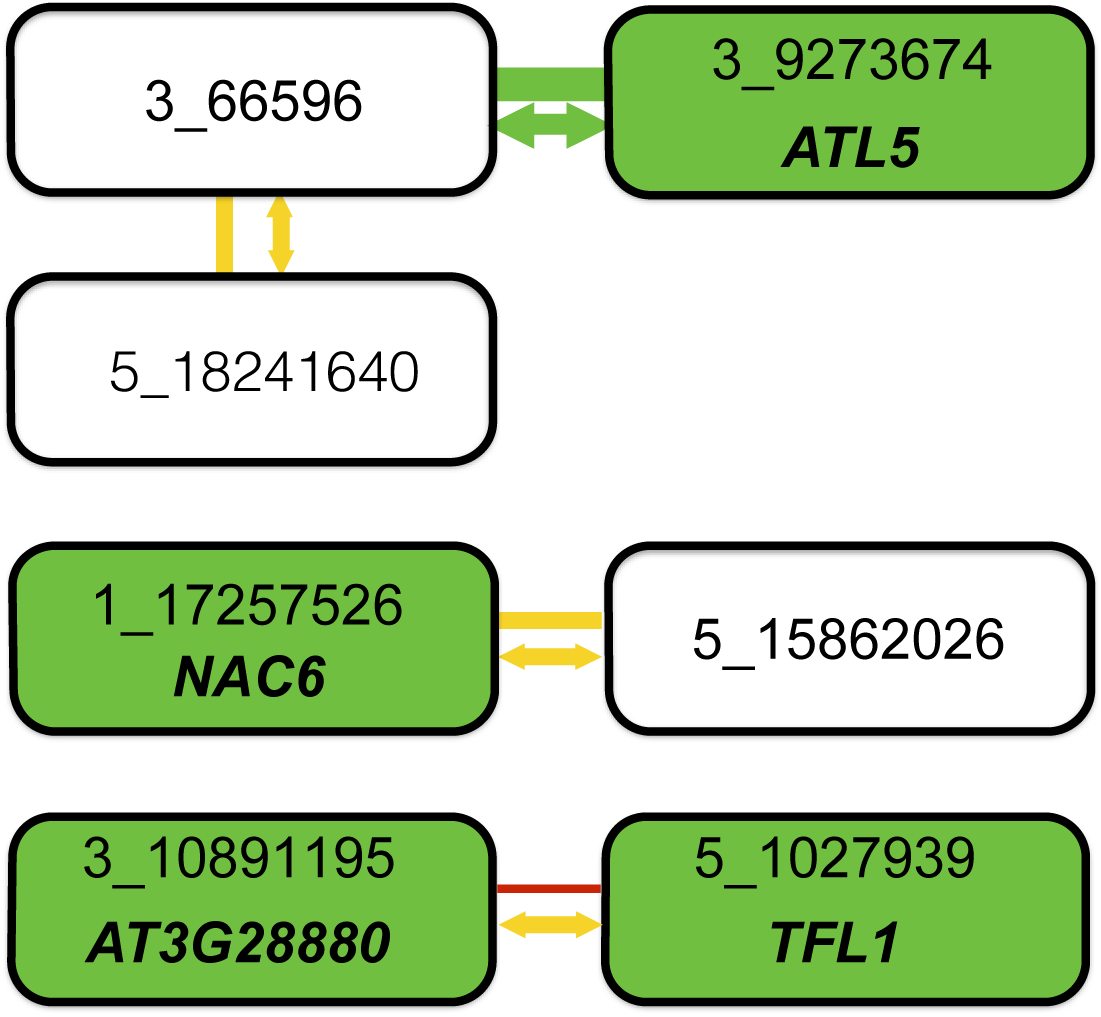
Summary of combined statistical and functional support for loci underlying root length. Four epistatic pairs, involving seven unique loci (connected boxes), were identified at a genome-wide significance-threshold in a two-dimensional genome-wide GWA scan. Green, yellow, and red lines connect pairs of loci with very low (p = 0.003), intermediate (0.28 < p < 0.37), and high (p = 0.95) risk for the interaction being a false-positive when accounting for population size. The risk of the statistical epistatic association resulting from high-order LD to an unobserved functional variant in the genome (i.e. “apparent epistasis”) is illustrated by arrow color, in which yellow indicates an intermediate risk (0.18 < p < 0.42) and green a very low risk (p = 0.0021). Green boxes indicate loci that the T-DNA insertion line analyses suggest the named genes to be involved in root development. When considering the joint statistical and functional results, two pairs emerge as highly likely true positive two-locus associations: 3_66596/3_9273674 due to very strong statistical support and one identified functional candidate gene, and 3_10891195/5_1027939 where the identification of functional candidate genes at both loci suggest that the two-locus association in the original genome-wide scan is true despite the lower statistical support in after the conservative statistical correction for sample-size. For the other two pairs, the results are inconclusive. There is strong support for one of the two associated loci (3_66596 from its statistical interaction with 3_9273674 and 1_17257526 by the detection of the a functional candidate gene in the T-DNA analysis), but weaker support for the second locus. Further work is thus needed to conclude whether these pairs represent true positive two-locus associations, or whether they are false-positives due to the small population-size or high-order LD (“apparent epistasis”) to unknown functional variants.

#### The previously uncharacterized gene At3g28880 contributes to root length

A locus on chromosome 3 (10,891,195 - SNP 3_10891195; Table 2) was identified due to its statistical epistatic interaction with a locus on chromosome 5 (1,027,939 bp - SNP 5_1027939; Table 2). SNPs in LD with 3_10891195 were found in three genes: At3g28865, At3g28870, and At3g28880 (Table 5). None of these genes previously had been associated with root length. By examining available insertion mutants, we established a novel role for the gene At3g28880 in root development; its mutant showed significantly decreased mean root length compared to wild-type controls (p = 4.5 × 10^−10^; Fig 4), whereas At3g28865 did not. At3g28880 encodes a previously uncharacterized ankyrin family protein. There is no information on developmental expression patterns for this gene; although the gene’s putative regulatory region is accessible in seeds and seedling tissue [44]. Across the tested accessions, a number of promoter-proximal SNPs were found in At3g28880, possibly altering gene regulation.

#### Mutant TFL1 increases mean root length

The SNP 5_1027939 was detected via its epistatic interaction with the At3g28880 locus on chromosome 3 (Table 2). This polymorphism and the only other polymorphism in LD are both intergenic between the genes At5g03840 and At5g03850 (Table 5). Phenotyping the insertion mutants for these genes revealed that the mutant for At5g03840, which encodes *TFL1*, showed significantly longer roots than wild-type controls (p = 5.7 × 10^−10^; Tukey post-hoc test; Fig 4), whereas no significant phenotypic effect was found for the At5g03850 mutant. Interestingly, these results implicate *TFL1* as a repressor of root growth.

*TFL1*’s role in floral initiation and morphology are well-established in many plant species [47]. It also plays a role in regulating protein storage in vegetative tissues, such as roots or seeds [48]. *TFL1* is highly expressed in both adult and seedling roots [45]. Our finding that *TFL1* controls root length is particularly poignant as a previous study reported no root length defects for a different *tfl1* mutant with multiple other pleiotropic phenotypes [49]. Our finding extends the functional reach of this multi-functional regulator.

## Discussion

This study was designed to identify genes regulating seedling root length. The narrow-sense heritability for this trait was not significantly different from zero in our population, whereas there was significant broad-sense heritability (Table 1). Notably, we observed a very large contribution of epistatic variance to the phenotypic variation in mean root length. With these findings in hand, we completed a comprehensive set of GWA analyses to identify additive and epistatic loci affecting this trait.

Consistent with the low narrow-sense heritability for mean root length, we were unable to identify any additive loci that reached the Bonferroni-corrected significance threshold in our analyses. By accounting for the contribution of pairwise epistatic interactions to the genetic variance of root length, however, we identified seven loci involved in four unique epistatically interacting pairs of loci that together explained a large fraction of the phenotypic variance in the population (Table 2). One of these pairs is highly significant also when accounting for the small population-size, whereas there is a risk that two of the three remaining pairs are false-positives due to this. We also showed that it is unlikely that the statistical epistatic interaction for the most significant pair would result from “apparent epistasis” due to a major, hidden variant in the genome. This risk is, however, not negligible for the three less significant pairs. Based on this, we conclude that despite the fact that much of the genetic variance present in this population can be explained by the genome-wide significant pairwise epistasis detected amongst these loci, additional evidence is needed before any firm conclusions can be drawn based on the other three epistatic pairs detected in the GWA analysis.

All four pairs detected in the initial epistatic GWA display an epistatic cancellation of additive genetic variance (Fig 2c). The presence of this type of genetic architecture for root-length in this population explains the low narrow-sense heritability in the population and why no loci could be found in the standard GWA analyses. Although further studies are needed to explore how general this phenomenon is, our result contrasts the common view in quantitative genetics that most genetic variance for complex traits is expected to be additive [5]. It further illustrates the importance of taking a richer modeling approach in GWA analyses to improve the sensitivity as a too strict focus on additive effects in the initial statistical analyses might lead to important findings being missed and lead to an underestimation of the full adaptive potential due to the epistatic suppression of selectable additive genetic variance [10–12].

Our finding of extensive statistical epistasis in this study should not be interpreted as evidence for root development being a biological trait with exceptionally high levels of biological interactions. Rather it is important to notice that, although the levels of statistical epistasis detected here originate in the epistatic genotype-phenotype maps for these pairs, the exceptionally high levels of epistatic variance in the population is even more due to the allele-frequencies for the involved loci in the population (Fig 2c). Hence if the loci displaying the G-P map in Fig 2c instead, for example, would be present at intermediary and equal frequencies, they would contribute high levels of additive genetic variance instead. Consequently, our results is an important reinforcement of the fact that the levels of epistatic variance for a trait in a population will be a dynamic outcome of changes in genotype-frequencies over time even though the underlying genotype-phenotype map might be constant [7–13].

In practice, however, neither allele frequencies nor genotype-phenotype maps will be known for the loci contributing to a particular trait. As illustrated by our findings for root length in this population, it is important to utilize GWA analysis methods that account for both additive and non-additive genetic variance to detect the contributing loci. Once this is done, further theoretical and experimental work is needed to identify the underlying genotype-phenotype maps and dissect the molecular underpinnings of the statistical associations regardless of whether these were detected mainly via their contributions to the additive and/or epistatic genetic variance. We here propose how to statistically quantify the risks of each inferred epistatic GWA association to guide further experimental work based on these findings. We find that there were statistical reasons to interpret three of the detected epistatic pairs with caution, both due to an increased risk of false-positives due to the small population-size and an inflated risk for “apparent epistasis”. Three novel functional candidate genes were, however, identified in loci detected as part of these associations. This illustrates clearly that the ultimate contribution from a study will rely on the combined use of appropriate statistical analyses to quantify the confidence in a particular locus and an in-depth experimental dissection of the detected associations before a conclusion can be reached about which loci contribute to a particular trait.

Functional dissection of epistasis, however, is a daunting task in natural populations and therefore model organisms are an indispensable resource in this work. A first step in taking on this challenge is to reduce the list of potential candidate genes and mutations for further, in-depth functional explorations. Here, we use mutational analysis of the genes that harbor SNPs in LD with the associated SNPs in the epistatic GWA analysis to identify the most likely candidate genes for the inferred associations. We tested six of the seven epistatic regions with available T-DNA insertion mutants. For four of the loci, we succeeded in identifying candidate genes affecting root length, three of which were newly implicated in root development. The majority of single-gene mutations affects do not affect root length [4, 35, 50]. Our results demonstrate the benefits of using statistical epistatic analyses in combination with experimental analysis of T-DNA insertion mutants to identify novel functional candidate genes involved in root development and provide a basis for future detailed studies of the complex gene networks involved in this trait (Fig 4).

It will be a major endeavor to functionally explore the biological mechanisms contributing to the statistical epistatic associations. None of the genes affecting root development contain non-synonymous mutations with likely phenotypic effects. Also, none of these genes have previously been described to interact with their respective epistatic locus. Future functional studies will have to explore the effects of the observed regulatory SNPs in transgenic plants in several backgrounds to study the whether these statistical interactions are the result of an underlying biological interaction or not. This endeavor will require significant time and resources, yet we believe that *A. thaliana* root length represents an excellent model trait for future in-depth studies of this type of complex signals arising from GWA studies.

## Materials and Methods

### Phenotyping

In order to reduce environmental variation among accessions, eighteen individuals from each of the 93 accessions studied in Atwell et al (2010) (Table S1) were vernalized at 4° as seedlings for six weeks to synchronize growth and flowering. The five most developmentally advanced seedlings from each accession were then transferred to soil in a randomized design. Plants were grown in long-day conditions at 22°. Flats were rotated three times per week to reduce position effects on plant development. Seeds were collected over a period of three months as the plants dried. Equal numbers of seeds were pooled from 3-5 parent plants to reduce the environmental contribution of plant placement across flats of plants. Ethanol sterilized seeds were planted on 1× Murashige and Skoog (MS) basal salt medium supplemented with 1× MS vitamins, 0.05% MES (wt/vol), and 0.3% (wt/vol) phytagel in a semi-randomized design with n = 70 per accession.

Four sets of twenty-three accessions plus a control accession (Col-0) made up a set. Each set was replicated three times providing the standard error for variance, with a total n = 210 planted for each accession. The seeds were stratified at 4° for three days and then grown for seven days in darkness with the plates in a vertical position. A photograph of each plate was taken, and root length was measured manually using the ‘freehand’ function in ImageJ1.46r [51]. Non-germination, missing organs, and delayed development were noted.

### Data standardization

Thirty-one seedlings with hypocotyls less than 5mm were removed because it is likely that germination was severely delayed [34]. Between-set differences were corrected by subtracting the difference between Col-0 in each replicate and set and the global mean for Col-0. Systematic differences between replicates were still present and corrected within each set by using the residuals from a model in which root length was explained by the within-set replicates.

### Heritability estimation

Heritability of root length variance and mean were estimated using the repeated measures with genotype as a fixed effect using ANOVA. To parse the contribution of additive and epistatic effects, a linear mixed model including both additive and epistatic effects as random effects was fitted using the R/hglm package [37], i.e.

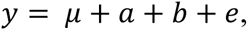

where

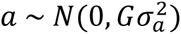

are the individual-level additive genetic effects, and G is the genomic kinship matrix;

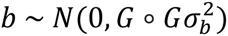

are the individual-level epistatic effects, and G is the genomic kinship matrix; *e* are independent and normally distributed residuals. The narrow sense heritability was estimated as the ratio of the additive genetic variance to the total phenotypic variance, and the broad sense heritability was estimated as the ratio of the sum of the additive and epistatic variance to the total phenotypic variance.

### Genome-wide association analysis

Using the transformed mean and the genotypes for the 93 accessions from Atwell et al. (2010), we performed a series of analyses to detect marker-trait associations. For the standard additive GWA analysis with population structure controlled, we used the function egscore() from R/GenABEL [38]. The functions bigRR() and bigRR_update() in the R/bigRR package [39] were used to perform a GWA analysis, in which all genome-wide SNP effects are modeled simultaneously as random effects, and the effects were estimated via the generalized ridge regression method HEM. The maximum absolute effect sizes in 1000 permutations were extracted to determine an empirical 5% genome-wide significance threshold (p = 0.0020).

To screen the genome for pairs of epistatic SNPs, we used the PLINK –epistasis procedure [40] that is based on the model:

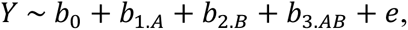

which considers allele by allele epistasis in. In the analysis, all possible pairs of SNPs with a minor allele frequency > 25% were tested.

We filtered out the pairs where there were fewer than four accessions in the minor two-locus genotype-class. Further, all combinations in which the p-value was > 1 × 10^−8^ were also removed as it was considered unlikely that they would be significant after correction for population structure and applying a multiple-testing corrected significance threshold. For the remaining pairs, a linear mixed model including fixed, additive and epistatic effects, as in the PLINK based initial scan, were fitted together with a kinship correction for population stratification, as in the single locus analyses, using the package R/hglm. We derived a multiple-testing corrected significance threshold for this epistatic analysis (p = 3.2 × 10^−10^) by estimating the number of independent tests based on the number of estimated LD blocks in the genome. For this, we used the fact that the *A. thaliana* genome is approximately 125Mb with average LD-blocks of about 10kb (12,500; [41]), and then applied a Bonferroni correction for the 78 million independent tests performed. Seventy-eight million double-recessive pseudo markers were created, and the R/p.exact package [42] was used to double-assess the exact false discovery rate of each detected epistatic pair based on its estimated epistatic effect size and allele frequency. Using data on the sixty-three accessions from the GWAs that were unambiguously identified in the 1001 Genomes data [22], we then identified additional SNPs in LD with the leading SNP using the function *LD* in the *genetics* package in R across a 10kb region around the marker.

### High-order LD between epistatic pseudo markers and whole-genome sequencing variants

From 728 whole-genome sequences in *A. thaliana* 1001 Genomes Project (http://www.1001genomes.org), we extracted all the sequencing variants. All four significant pseudo-markers had at least four accessions with the double-recessive genotype (i.e. the minor allele for the epistatic pseudo-marker) in the 728 sequenced accessions. For each of the pseudo-markers, we therefore sampled 93 accessions from the 728 sequenced ones, keeping the same allele frequency as for the pseudo-marker representing the epistatic pair detected in the GWA dataset, and obtained the genome-wide, high-order LD *r*^2^ distribution across all the sequenced variants. This procedure was repeated 50 times for each pseudo-marker, and we calculated the estimates and standard errors of the *r*^2_mean_^, *r*^2_max_^ and estimated the risk for “apparent epistasis” as the probability of observing an *r*^2^max > 0.8 in a particular random sample.

### Examination of SNP-associated genes

The Ensembl Variant Effect Predictor based on the TAIR10 release of the *A. thaliana* genome was used to determine the effects of the leading SNPs and the SNPs in high LD with them. Genes were considered as candidates if they were within 1kb of a variant. Expression of the candidate genes was determined using the BAR eFP Arabidopsis browser [45].

### Validation of interacting loci

T-DNA lines were obtained for the candidate genes (Table S2). Root length was ascertained as described above (n = 20). Tukey’s HSD post hoc test was used to compare root lengths between the T-DNA lines and the wild-type accession (*Col-0*).

## Acknowledgments

We thank Mats Pettersson and Ronnie Nelson for useful discussions and Karla Schultz and members of the Queitsch lab for experimental assistance.

## Supporting information captions

**Table S1. *A. thaliana* accessions phenotyped for GWA.**

**Table S2. T-DNA lines tested for root phenotypes.**

**Fig S1. No significant associations are found in GWAS for root length using an additive model.**

(A) In the GWAS based on an additive model no SNPs reached the Bonferroni threshold. (B) Using a whole-genome generalized ridge regression model, in which SNPs were modeled as random effects, no SNPs reached the Bonferroni threshold.

**Fig S2. Similar genotype-phenotype maps for all significantly interacting pairs of loci.**

(A-C) Mean root length is displayed for all four two-locus genotype-classes for the four pairs of significantly interacting loci. The major allele is indicated by -1 and the minor allele is indicated by 1. The combination of two minor alleles (1,1) always has the longest root length compared to the other three-allele combinations, which are more similar to one another. Pictured are mean root lengths for the epistatic pairs for SNPs (A) 3_66596 and 3_9272294/3_9273674, (B) 3_66596 and 5_18241640, and (C) 3_10891195 and 5_1027939.

